# A Neurodevelopmental Disorder Caused by Mutations in the VPS51 Subunit of the GARP and EARP Complexes

**DOI:** 10.1101/409441

**Authors:** David C. Gershlick, Morié Ishida, Julie R. Jones, Allison Bellomo, Juan S. Bonifacino, David B. Everman

## Abstract

GARP and EARP are related heterotetrameric protein complexes that associate with the cytosolic face of the *trans*-Golgi network and recycling endosomes, respectively. At these locations, GARP and EARP function to promote the fusion of endosome-derived transport carriers with their corresponding compartments. GARP and EARP share three subunits, VPS51, VPS52 and VPS53, and each has an additional complex-specific subunit, VPS54 or VPS50, respectively. The role of these complexes in human physiology, however, remains poorly understood. By exome sequencing, we have identified compound heterozygous mutations in the gene encoding the shared GARP/EARP subunit VPS51 in a six-year-old patient with severe global developmental delay, microcephaly, hypotonia, epilepsy, cortical vision impairment, pontocerebellar abnormalities, failure to thrive, liver dysfunction, lower extremity edema and dysmorphic features. The mutation in one allele causes a frameshift that produces a longer but highly unstable protein that is degraded by the proteasome. In contrast, the other mutant allele produces a protein with a single amino-acid substitution that is stable but assembles less efficiently with the other GARP/EARP subunits. Consequently, skin fibroblasts from the patient have reduced levels of fully-assembled GARP and EARP complexes. Likely because of this deficiency, the patient’s fibroblasts display altered distribution of the cation-independent mannose 6-phosphate receptor, which normally sorts acid hydrolases to lysosomes. Furthermore, a fraction of the patient’s fibroblasts exhibit swelling of lysosomes. These findings thus identify a novel genetic locus for a neurodevelopmental disorder and highlight the critical importance of GARP/EARP function in cellular and organismal physiology.

## 1 Introduction

The “<u>G</u>olgi-<u>A</u>ssociated <u>R</u>etrograde <u>P</u>rotein” (GARP) and “<u>E</u>ndosome-<u>A</u>ssociated <u>R</u>ecycling <u>P</u>rotein” (EARP) complexes are structurally related, four-subunit complexes that promote SNARE-dependent fusion of endosome-derived transport carriers with target organelles in most eukaryotic cells^1,2^. GARP and EARP comprise three shared subunits named VPS51, VPS52 and VPS53, and one complex-specific subunit, either VPS54 (GARP) or VPS50 (EARP)^3,4,5,6,7,8^ (Fig. 1A). GARP is mainly associated with the cytosolic face of the *trans*-Golgi network (TGN), where it participates in the tethering and fusion of tubular-vesicular transport carriers derived from endosomes, as part of a process known as “retrograde transport”^9,10,11^ (Fig. 1B).

**Figure 1:**
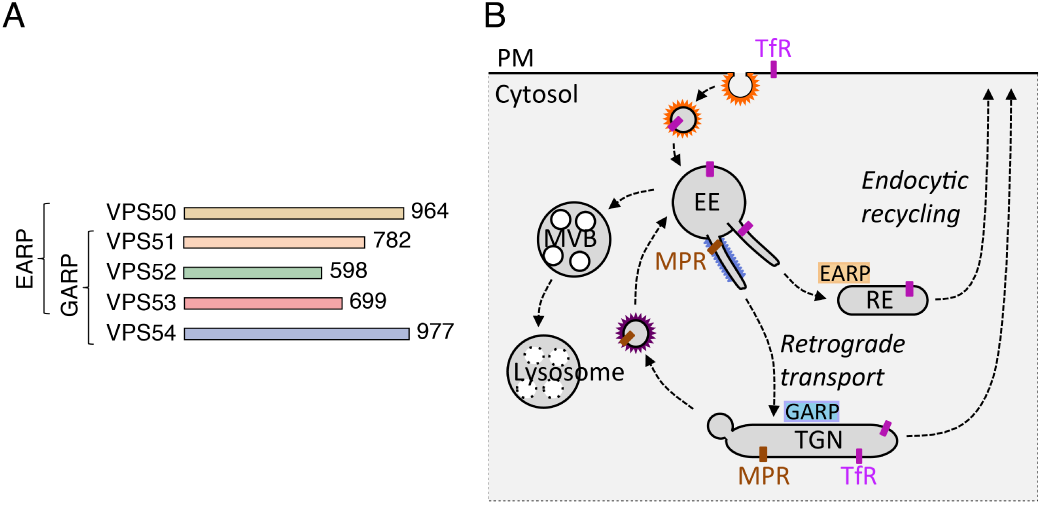
Characteristics of GARP and EARP. (A) The GARP complex is composed of VPS51, VPS52, VPS53 and VPS54 subunits, whereas the EARP complex is composed of VPS50, VPS51, VPS52 and VPS53 subunits^1,2^. GARP and EARP were also previously referred to as VFT (VPS fifty three)^3^and GARPII^7^, respectively. VPS51 is also known as Ang2 (another new gene 2)^6,38^, and VPS50 as syndetin^8^ or VPS54L^7^. The number of amino acids in each protein is indicated (according to UniProt, http://www.uniprot.org/). The subunits of GARP and EARP are elongated proteins belonging to the family of “complexes associated with tethering containing helical rods” (CATCHR)^39^. The four subunits of the mammalian complexes are thought to assemble via their N-terminal regions into an X-shaped structure^40^. (B) Schematic representation of the localization and function of GARP and EARP (adapted from ref. 8). GARP mediates tethering and fusion of endosome-derived carriers to the TGN, while EARP functions in tethering and fusion of endosome-derived carriers with recycling endosomes. GARP also plays a minor role in this latter pathway. EE: early endosome. MVB: multivesicular body. PM: plasma membrane. RE: recycling endosome

Examples of mammalian proteins that undergo GARP-dependent retrograde transport are the cation-independent mannose 6-phosphate receptor (CI-MPR) that cycles between the TGN and endosomes in the process of sorting acid hydrolase precursors to lysosomes^12^ and the vesicle SNARE VAMP4 that is involved in fusion of endosome-derived carriers with the TGN^11^. GARP is also required for retrograde transport of sphingolipids from endosomes to the TGN^13^. EARP, on the other hand, is mainly associated with the cytosolic face of endosomes marked by the small GTPase Rab4, and promotes the recycling of proteins such as the transferrin receptor (TfR) from endosomes to the plasma membrane^7,8^ (Fig. 1B). Studies in the nematode *C. elegans* have additionally implicated EARP in cargo sorting to dense-core vesicles in neurons and neuroendocrine cells^14^. Finally, the VPS51 subunit of GARP/EARP was shown to be defective in *fat-free* (*ffr*) mutant zebrafish, which display abnormal Golgi structure and defective lipid transport in the intestinal tract^15^.

The functions of GARP and EARP in intracellular transport are essential for the viability of mammalian organisms, as demonstrated by the embryonic lethality of mice with homozygous null mutations in subunits of these complexes^16,17,18,19^ (www.informatics.jax.org). In addition, a spontaneous, recessive mutation (A>T nucleotide 72 of exon 23) causing substitution of glutamine for leucine-967 (L967Q) in the GARP-specific VPS54 subunit is responsible for the phenotype of the wobbler (*wr*) mouse^17^. This substitution causes structural destabilization and partial degradation of VPS54, resulting in reduced levels of the fully-assembled GARP complex^20^. Wobbler mice are viable but develop motor neuron degeneration similar to that of amyotrophic lateral sclerosis (ALS)^21^. Despite the similarities between the wobbler phenotype and ALS, however, mutations in VPS54 have not been shown to cause ALS^22^. Recent genome-wide linkage analysis in conjunction with whole exome sequencing identified compound heterozygous mutations (c.2084A>G and c.1556+5G>A) in the gene encoding the shared VPS53 subunit of GARP and EARP in patients with a condition known as progressive cerebello-cerebral atrophy type 2 (PCCA2) or pontocerebellar hypoplasia type 2E (PCH2E) (OMIM #615851), a congenital disorder characterized by intellectual disability, progressive microcephaly, spasticity and epilepsy^23^. These mutations are predicted to affect the C-terminal region of VPS53^23^, although the effects of these mutations on the properties of the protein were not experimentally addressed. Phenotypic overlap between PCCA2 and an autosomal recessive condition known as progressive encephalopathy with edema, hypsarrhythmia and optic atrophy (PEHO)^24^ has also been described, as two siblings with a PEHO-like syndrome were recently found to have compound heterozygous mutations in VPS53^25^. This disorder was primarily characterized by severe developmental delay, postnatal microcephaly, progressive cerebellar and cerebral atrophy, seizures, ophthalmologic abnormalities, facial/limb edema, and dysmorphic features^25^.

Herein we report the identification of compound heterozygous mutations (c.1468C>T and c.2232delC) in the gene encoding the GARP/EARP subunit VPS51 in a 6-year-old patient with severe global developmental delay, pontocerebellar abnormalities, microcephaly, hypotonia, epilepsy, cortical vision impairment, failure to thrive, liver dysfunction, lower extremity edema, and dysmorphic features. Biochemical analyses show that a frameshift caused by the c.2232delC mutation (allele 1) produces a longer but highly unstable protein. On the other hand, the c.1468C>T mutation (allele 2) produces a protein with a single amino acid substitution (R490C) that is stable but assembles less efficiently with the other GARP/EARP subunits. As a consequence, skin fibroblasts from the patient have reduced levels of fully-assembled GARP and EARP complexes, and display altered distribution of the CI-MPR and swelling of lysosomes. These findings thus identify a new genetic locus for a complex neurodevelopmental disorder characterized by pontocerebellar abnormalities and other systemic manifestations, and highlight the critical importance of GARP/EARP function in cellular and organismal physiology.

## 2 Results

### 2.1 Clinical phenotype

The patient is a 6-year-old Caucasian female who presented in early infancy with cholestatic hepatitis, feeding difficulties, and failure to thrive. Her subsequent clinical course has included ongoing liver dysfunction requiring treatment with the bile acid ursodiol, gastrostomy tube dependence, hypotonia, marked microcephaly, severe global developmental delay with an inability to sit and a lack of spoken language, electrical status epilepticus in sleep (ESES) requiring anticonvulsant therapy, cortical vision impairment with associated strabismus, sleep apnea, a gastric volvulus, constipation, asthma, and episodic respiratory infections including RSV, adenoviral pneumonia, and a pleural effusion.

Multiple physical examinations have shown minor dysmorphic features, including epicanthal folds, long eyelashes, slightly overfolded ears, an upturned nasal tip, a thin upper lip, a high-/narrow anterior palate, full/rounded cheeks, a low posterior hairline, single flexion creases on the fifth fingers, mild clubbing of the thumbnails and other fingernails, and increased hair on the upper back. At 6 years of age, she was also noted to have bilateral lower extremity edema. A brain MRI at 4 months of age was normal, while repeat MRI scans at 4 and 6 years of age (Fig. 2A) showed multiple abnormalities, including a small cerebellar vermis, a small pons/brainstem, an enlarged infra-vermian cistern with suspicion for a Dandy-Walker variant, a small hippocampus, a thin corpus callosum, and abnormal white matter signal in the cerebral hemispheres. A more detailed clinical report is included in Supplemental Data.

**Figure 2:**
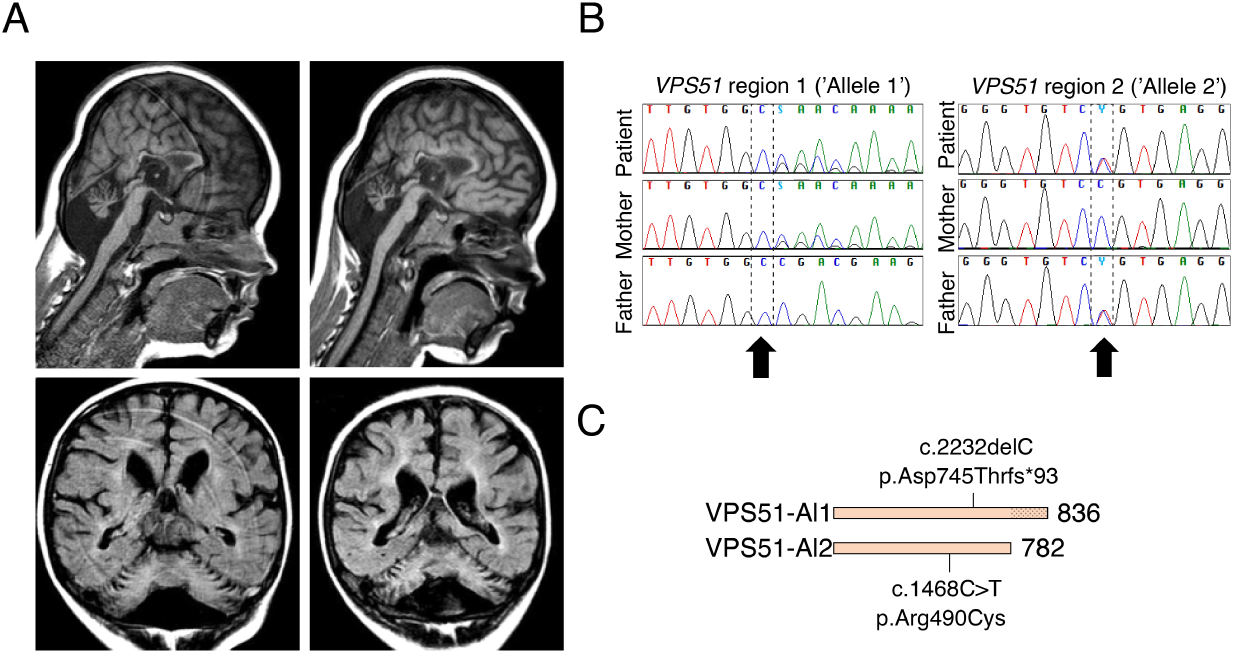
Clinical and sequencing findings in the patient. (A) Selected brain MRI images from ages 4 years 6 months (left) and 6 years 3 months (right). Note pontocerebellar hypoplasia, prominent infra-vermian cistern, and hypoplastic corpus callosum. (B) Sanger sequencing of DNA samples from the patient and both parents confirming mutations in *VPS51* and compound heterozygosity, as originally found by whole-exome sequencing (see Fig. S1). (C) Representation of mutations in VPS51. The mutation in allele 1 causes replacement of threonine for aspartate-745 and a frameshift that adds 54 random amino acids at the C-terminus of the protein. The mutation in allele 2 results in substitution of cysteine for arginine-490.

Biochemical testing has been notable for abnormalities suggesting a possible congenital disorder of glycosylation (CDG), including evidence of hypoglycosylation on serum transferrin isoelectric focusing (manifested by a slightly low percentage of trisialotransferrin and slightly increased percentages of asialo- and monosialo-transferrin) as well as an abnormal pattern of N-and O-glycosylation. Other biochemical tests were normal or non-diagnostic (see Supplemental Data).

### 2.2 Identification of compound heterozygous variants in *VPS51*

Whole exome sequencing was performed through the Molecular Diagnostic Laboratory at the Greenwood Genetic Center using genomic DNA from the patient and both parents and identified compound heterozygous variants in *VPS51* (reference sequence NM 013265.3) (Fig. 2B,C; Fig. S1), including a maternally inherited frameshift alteration (allele 1: c.2232delC; p.Asp745Thrfs*93) that was considered pathogenic and a paternally inherited missense alteration (allele 2: c.1468C>T; p.Arg490Cys) that was classified as a variant of uncertain significance (VUS) prior to the publication of the ACMG variant interpretation guidelines^26^.

The allele 1 variant results in the substitution of threonine for aspartate at amino acid 745 and a downstream frameshift that causes the normal stop codon to be skipped. The resulting open reading frame is 202 nucleotides longer and encodes a protein with an additional 54 extraneous amino acids C-terminal to aspartate 745 (Fig. 2C). This variant has not been reported in the Human Gene Mutation Database (HGMD) (http://www.hgmd.cf.ac.uk/ac/index.php). Allele 2 has been reported in the public SNP databases but limited information is available to support it as benign.

Whole exome sequencing also identified heterozygous VUSs in three other genes. These included a paternally inherited missense variant (c.398C>G; p.Pro133Arg) in *ALG1*, a maternally inherited synonymous variant (c.2022C>G; p.Val674Val) in *COG4*, and a maternally inherited intronic variant (c.2014 *-*4A>G) in *ALG13*. Pathogenic alterations in these genes have been associated with autosomal recessive CDG type Ik (OMIM #608540), autosomal recessive CDG type IIj (OMIM #613489), and X-linked recessive CDG type Is (OMIM #300884), respectively. The recessive nature of these disorders, however, made it unlikely that the monoallelic variants in *ALG1, COG4* and/or *ALG13* were the cause of the disease in our patient.

Reanalysis of the exome data was performed approximately 18 months after the original analysis and did not identify any other significant variants but did lead to reclassification of the *ALG13* variant as benign, based on 339 hemizygotes and 748 heterozygotes having been reported in the gnomAD database (http://gnomad.broadinstitute.org/). According to the ACMG guidelines on variant interpretation, a gene-disease association must be clinically validated prior to using the ACMG criteria; therefore, both of the *VPS51* variants would currently be classified as VUSs. Once the gene-disease association for *VPS51* has been validated, the use of ACMG criteria would classify the c.2232delC alteration as pathogenic and the c.1468C>T change as likely pathogenic^26^.

### 2.3 The allele 1 variant produces an unstable protein that is rapidly degraded by the proteasome

To assess the effects of the mutations on the properties of VPS51, we replicated them in a mammalian expression plasmid encoding human VPS51 tagged with either one copy of the green fluorescent protein (GFP) at the N-terminus or 13 copies of the myc epitope (13myc) at the C-terminus. WT and mutant forms of tagged VPS51 were expressed by transient transfection in HeLa cells.

Immunoprecipitation followed by immunoblotting of cells expressing GFP-tagged forms of normal VPS51 (GFP-VPS51) or allele 1 mutant VPS51 [GFP-VPS51(Al1)] showed that whereas the normal fusion protein ran at the expected molecular mass of ∼ 113 kDa, the mutant fusion protein was barely detectable (Fig. 3A). Furthermore, immunoblotting of extracts from cells expressing 13myc-tagged normal VPS51 (VPS51-13myc) or allele 1 mutant VPS51 [VPS51(Al1)-13myc] revealed that the mutant protein was expressed at lower levels than the normal protein (Fig. 3B).

**Figure 3:**
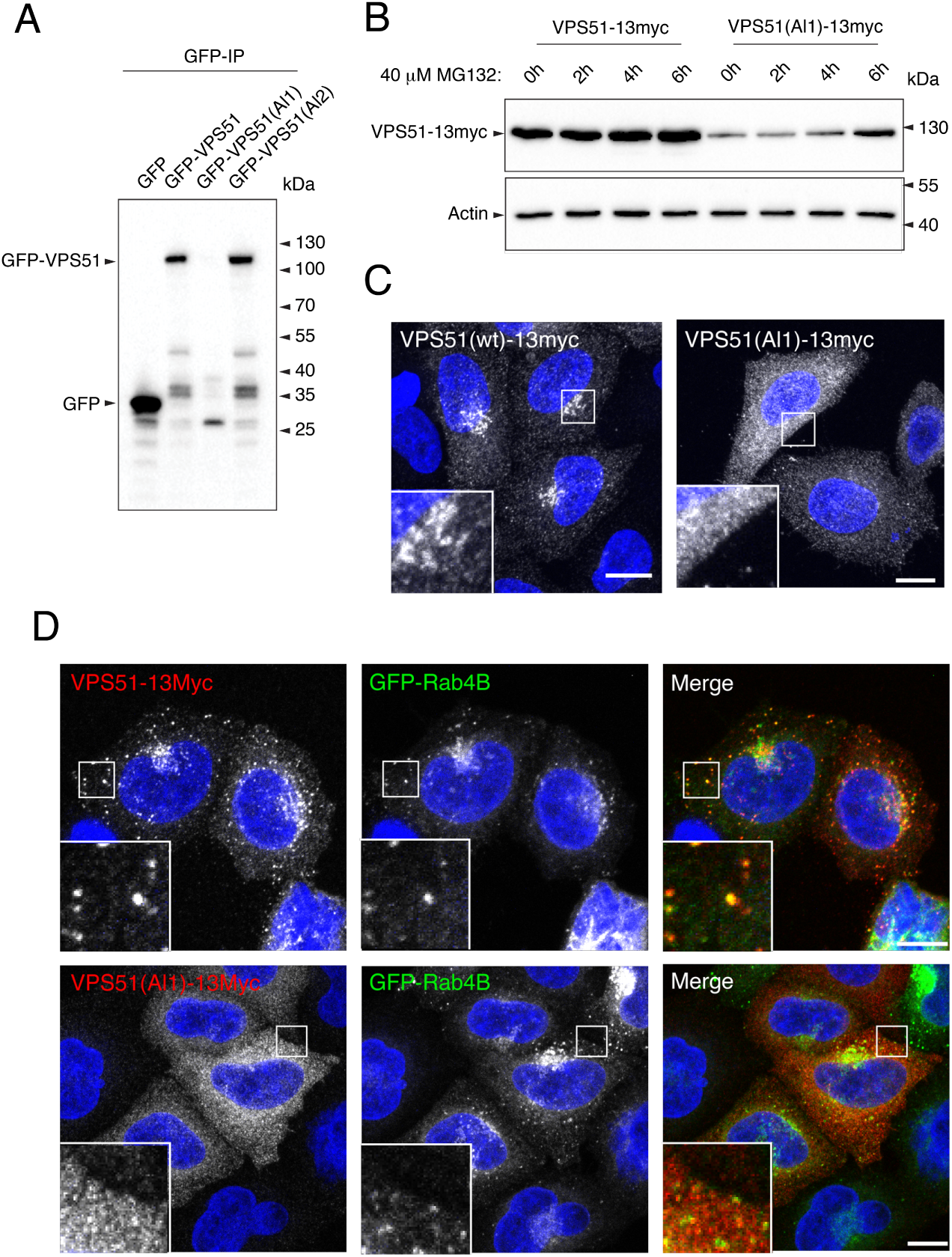
The VPS51(Al1) mutant protein is degraded by the proteasome and fails to associate with the TGN and endosomes. Figure 3: (A) Complementary DNAs encoding allele 1 and allele 2 mutant VPS51 tagged at the N-terminus with GFP [GFP-VPS51(Al1) and GFP-VPS51(Al2)] were expressed by transient transfection in HeLa cells. GFP and GFP-tagged normal VPS51 (GFP-VPS51) were similarly expressed as controls. Cell lysates were subjected to immunoprecipitation with GFP-nanobody-conjugated beads and immunoblot analysis with antibody to GFP. The positions of molecular mass markers are indicated to the right. Notice the undetectable levels of full-length GFP-VPS51(Al1) and the presence of lower molecular mass products of this protein. (B) Complementary DNAs encoding normal and allele 1 mutant VPS51 tagged at the C-terminus with 13 copies of the myc epitope [VPS51-13myc and VPS51(Al1)-13myc] were expressed by transient transfection in HeLa cells. Cells were incubated in 40 *μ*M of the proteasome inhibitor MG132 for the indicated times, lysed, normalized for protein abundance by the Bradford protein assay and analyzed by immunoblotting with an antibody to the myc epitope. The positions of molecular mass markers are indicated to the right. Notice the increase in VPS51(Al1)-13myc levels over the course of the experiment. (C) HeLa cells transiently transfected with plasmids encoding VPS51-13myc and VPS51(Al1)-13myc were immunostained for the myc epitope and imaged by confocal microscopy. Notice that normal VPS51-13myc localizes to the TGN, as previously reported^6^. In contrast, VPS51(Al1)-13myc shows cytosolic staining. (D) HeLa cells were transiently co-transfected with plasmids encoding GFP-Rab4B together with VPS51-13myc or VPS51(Al1)-13myc, immunostained for the myc epitope and imaged by confocal microscopy. Notice that GFP-Rab4B promoted recruitment of VPS51-13myc but not VPS51(Al1)-13myc to endosomes. In C and D, nuclei were stained with DAPI (blue). Single channels are shown in grayscale. In the merged images in D, myc staining is shown in red and GFP uorescence is shown in green. Scale bars: 10 *μ*m. Inset scale bars: 2 *μ*m.

Incubation with the proteasome inhibitor MG132 did not change the levels of normal VPS51-13myc but progressively increased the levels of mutant VPS51(Al1)-13myc over a 6 h period (Fig. 3B). These observations indicated that the allele 1 mutant protein was unstable and subject to degradation by the proteasome.

We also performed immunofluorescence microscopy of HeLa cells expressing normal VPS-51-13myc or mutant VPS-51(Al1)-13myc. We observed that the normal protein mainly localized to the TGN (Fig. 3C), as previously reported^6^. This localization is consistent with integration of the normal protein into the endogenous GARP complex. In contrast, expression of the mutant protein was detected in very few cells, and in these cells the protein was largely cytosolic (Fig. 3C). The small GTPase Rab4A was previously shown to promote recruitment of the EARP-specific VPS50 subunit (also known as syndetin) to a subpopulation of endosomes^7,8^. We observed that co-expression of GFP-Rab4B with VPS-51-13myc similarly resulted in co-localization of both proteins to discrete puncta distributed throughout the cytoplasm (Fig. 3D), consistent with the endosomal localization of EARP^7,8^. On the other hand, in the few cells that showed expression of VPS51(Al1)-13myc, the protein was cytosolic and did not localize to GFP-Rab4B-positive puncta (Fig. 3D).

Taken together, the above results indicate that VPS51 mutant allele 1 is expressed at very low levels due to its targeting for degradation by the proteasome. Moreover, even in those cells in which some expression is detected, the mutant protein is not recruited to sites of GARP and EARP localization. These defects likely result from misfolding caused by truncation of the normal sequence and addition of a random sequence C-terminal to amino acid 745.

### 2.4 The allele 2 mutation produces a stable protein that is not efficiently assembled into the GARP and EARP complexes

In contrast to the allele 1 mutation, the substitution of cysteine for arginine 490 in allele 2 did not affect the expression of the GFP-tagged mutant protein [GFP-VPS51(Al2)] relative to the normal protein (GFP-VPS51), as determined by immunoblot analysis of transfected HeLa cells (Figs. 3A and 4A). However, mutant GFP-VPS51(Al2) showed reduced co-immunoprecipitation with VPS50 and VPS53 relative to normal GFP-VPS51 (Fig. 4A), consistent with impaired assembly of the mutant protein into the GARP and EARP complexes. Co-immunoprecipitation with VPS54 was not tested because of the lack of a suitable antibody to this subunit. In agreement with the co-immunoprecipitation results, immunofluorescence microscopy of transfected HeLa cells showed decreased TGN localization of mutant VPS51-Al2-13myc relative to normal VPS51-13myc (Fig. 4B,C). When co-expressed with GFP-Rab4B, mutant VPS51(Al2)-13myc showed co-localization of both proteins to cytoplasmic puncta, but the number of puncta per cell was lower for mutant VPS51(Al2)-13myc relative to normal VPS51-13myc (Fig. 4D,E).

**Figure 4:**
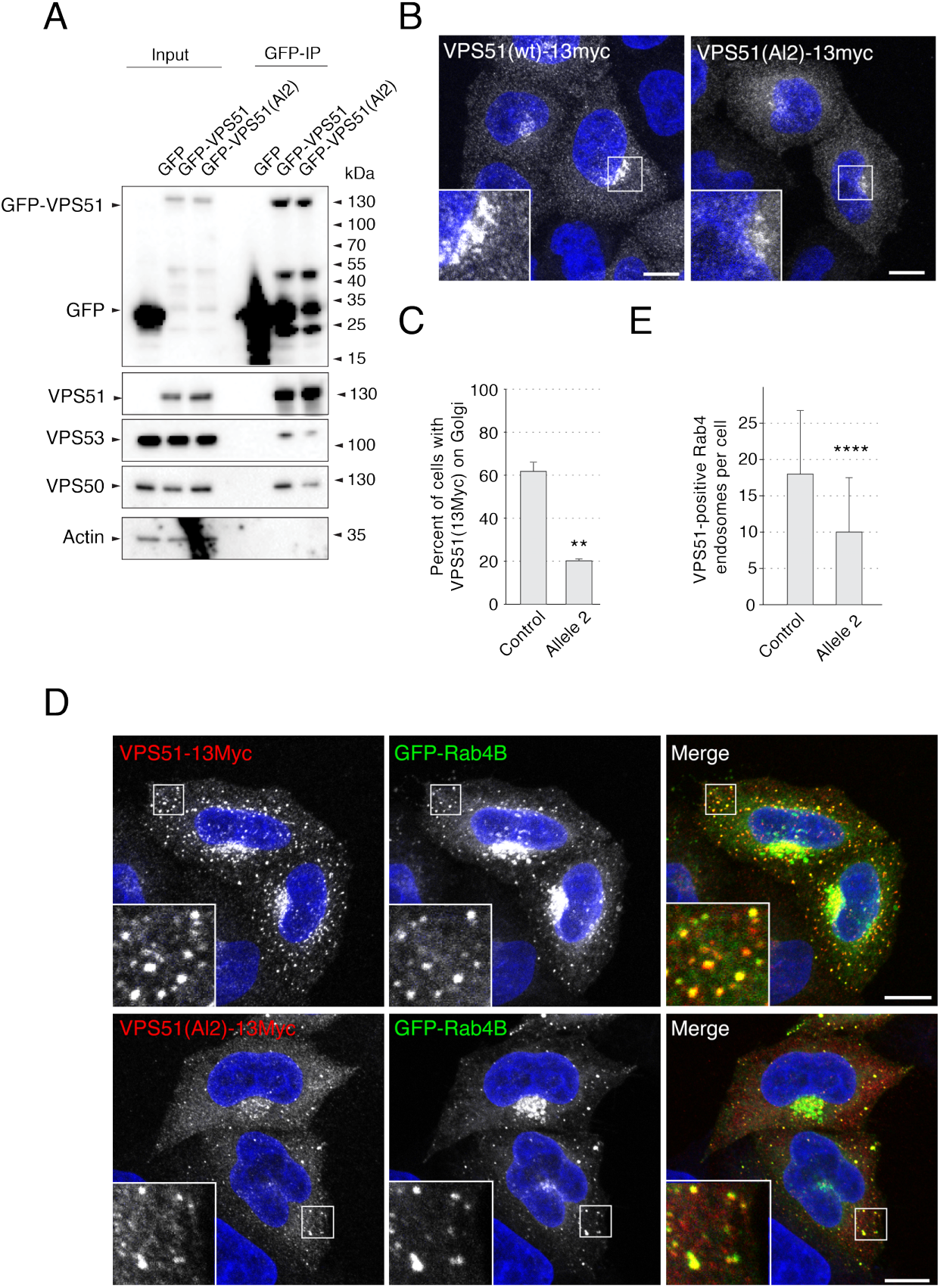
The VPS51(Al2) mutant protein is stable but assembles less efficiently into GARP and EARP. (A) Extracts from HeLa cells transiently transfected with plasmids encoding GFP, GFP-VPS51 or GFP-VPS51(Al2) were analyzed by pull down with a nanobody to GFP conjugated to magnetic beads, followed by SDS-PAGE and immunoblotting for GFP, endogenous GARP/EARP subunits and actin (loading and negative control). Positions of molecular mass markers are indicated on the right. Notice the similar levels of GFP-VPS51 and GFP-VPS51(Al2) and the decreased co-immunoprecipitation of VPS53 and VPS50 with GFP-VPS51(Al2). (B) HeLa cells were transiently transfected with plasmids encoding VPS51-13myc, immunostained for the myc epitope and imaged by confocal microscopy. (C) Quantification of the percentage of cells with VPS51-13myc staining at the TGN. Over 300 cells across 3 different independent experiments (over 100 cells per experiment per condition) were classified as either having residual TGN localization or no observable TGN localization. Percentages were calculated and compared with a one-tailed paired t-test, p-value = 0.0014. VPS51-13myc displayed a perinuclear (TGN) localization in over 60% of the transfected cells, whereas VPS51(Al2)-13myc showed a mostly cytosolic localization, with only ∼20% of cells having observable residual perinuclear staining. Moreover, VPS51(Al2)-13myc staining at the TGN was less-intense than that of VPS51-13myc in comparable cells. (D) HeLa cells were transiently co-transfected with plasmids encoding GFP-Rab4B together with VPS51-13myc or VPS51(Al1)-13myc, immunostained for the myc epitope and imaged by confocal microscopy. (E) Quantification of the number of GFP-Rab4B-positive endosomes that also contained VPS51-13myc or VPS51(Al1)-13myc. VPS51-positive Rab4B endosomes in 62 cells (wt VPS51) and 64 cells (allele 2), from images taken across 3 independent experiments were counted. Datasets were compared with a t-test, p-value = 5.5×10-7. Notice that GFP-Rab4B caused recruitment of VPS51-13myc. GFP-Rab4B also caused VPS51(Al2)-13myc recruitment to endosomes but the number of endosomes was smaller and there was more cytosolic background. In B and D, nuclear staining by DAPI is shown in blue. Single channels are shown in grayscale. In the merged images in D, myc staining is shown in red and GFP fluorescence is shown in green. Scale bars: 10 *μ*m. Inset scale bars: 2 *μ*m. Error bars = standard deviation.

From these experiments, we concluded that the allele 2 mutant VPS51 is stable but gets incorporated into the GARP and EARP complexes less efficiently, resulting in reduced levels of the fully-assembled complexes.

### 2.5 Decreased levels and membrane association of GARP and EARP in patient’s fibroblasts

We next examined the effects of the VPS51 mutations in skin fibroblasts from the patient. Immunoblot analyses of endogenous GARP/EARP subunits showed reduced levels of VPS51 (∼25%) as well as VPS50, VPS52 and VPS53 (all ∼50%) in fibroblasts from the patient relative to fibroblasts from a control individual (Fig. 5A,B).

**Figure 5:**
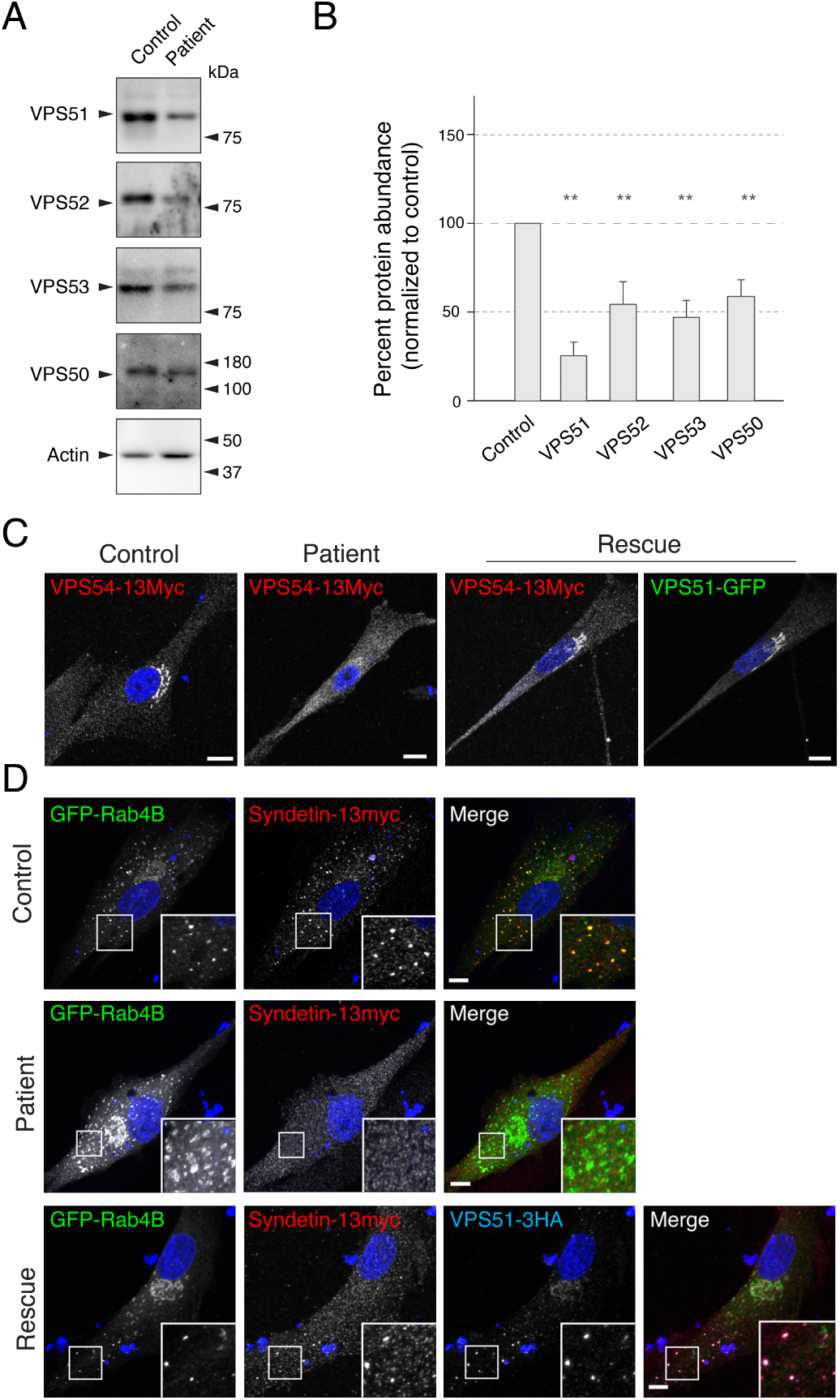
Patient’s fibroblasts show defects in GARP/EARP complexes. Figure 5: (A) Patient’s and control fibroblasts were lysed, protein abundance was normalized by the Bradford assay and samples were analyzed by SDS-PAGE and immunoblotting for endogenous GARP/EARP subunits and actin (negative control). Positions of the molecular mass markers are indicated on the right. (B) Quantification of fold differences in GARP/EARP subunit levels from three independent immunoblot assays. Actin loading control was used to normalize the expression. Samples were compared with a one-tailed paired t-test, p-values: VPS51 = 0.0097, VPS52 = 0.030, VPS53 = 0.0470, VPS50 = 0.0353. Notice the decreased levels of GARP/EARP subunits in the patient’s fibroblasts. C) Control and patient’s fibroblasts were transiently transfected with plasmids encoding VPS54-13myc (left two panels), and patient’s fibroblasts were transiently co-transfected with plasmids encoding VPS54-13myc and VPS51-GFP (right two panels). Notice that co-expression with VPS51-GFP recovers perinuclear localization of VPS54-13myc in the patient’s fibroblasts. (D) Control and patient’s cells were co-transfected with DNA plasmids encoding GFP-Rab4B and VPS50-13myc (top two panels), as well as VPS51-3HA (bottom panel). Notice that co-expression with VPS51-3HA recovers the localization of VPS50-13myc to the GFP-Rab4B-positive endosomes. In C and D, nuclear staining by DAPI is shown in blue. Single channels are shown in grayscale. In panel C, GFP signal is shown in green. In the merged images in D, GFP fluorescence is shown in green, myc staining in red, and, for the bottom panel, HA staining in blue. Scale bars: 10 *μ*m. Inset scale bars: 2 *μ*m. Error bars = standard deviation.

As none of the antibodies to GARP/EARP work for immunofluorescence microscopy of the endogenous proteins, we transfected the fibroblasts with plasmids encoding 13myc-tagged VPS54 (VPS54-13myc), which serves as a surrogate for GARP. We observed that VPS54-13myc exhibited localization to the TGN in control fibroblasts, as previously reported for other cell types^12^, and diffuse cytosolic localization in the patient’s fibroblasts (Fig. 5C).

We also co-expressed 13myc-tagged VPS50 (VPS50-13myc) (to label EARP) with GFP-Rab4B, and found that both proteins co-localized on punctate structures distributed throughout the cytoplasm in control fibroblasts (Fig. 5D), consistent with the endosomal localization of EARP^7,8^. In contrast, VPS50-13myc failed to localize to GFP-Rab4B puncta and was largely cytosolic in the patient’s fibroblasts (Fig. 5D). Both the localization of VPS54-13myc to the TGN and VPS50-13myc to Rab-4B-positive endosomes was restored by transfection of the patient’s fibroblasts with plasmids expressing normal VPS51 tagged with either GFP (VPS-51-GFP) or 3 copies of the HA epitope (VPS51-3HA), respectively (Fig. 5C,D). These experiments thus demonstrated that the mutations in VPS51 resulted in reduced levels of GARP and EARP and impaired association of both complexes with their corresponding intracellular compartments in the patient’s fibroblasts.

### 2.6 Altered distribution of the CI-MPR and abnormal lysosomes in patient’s fibroblasts

Previous studies showed that knock down (KD) of GARP subunits in HeLa cells altered the intracellular distribution of the CI-MPR^6,12,27^. In line with these findings, immunofluorescence microscopy showed that the CI-MPR exhibited bright juxtanuclear staining in control fibroblasts and faint, more dispersed cytoplasmic staining in the patient’s fibroblasts (Fig. 6A,B). In addition, we observed that the patient’s fibroblasts had lower levels of CI-MPR protein relative to the control fibroblasts, as analyzed by immunoblotting (Fig. 6C,F). These results are consistent with reduced retrieval to the TGN and increased turnover of the CI-MPR in the patient’s fibroblasts.

**Figure 6:**
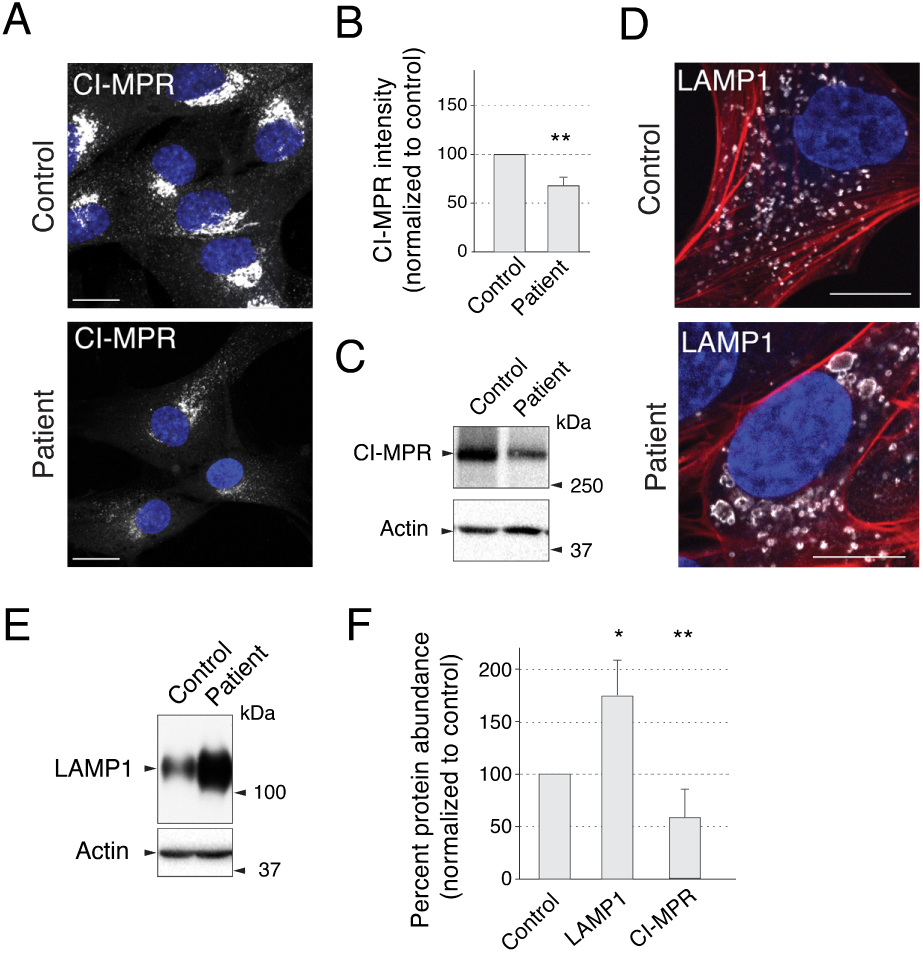
Patient’s fibroblasts show defects in GARP/EARP complexes. Figure 6: (A) Control and patient’s fibroblasts were immunostained with an antibody to endogenous CI-MPR. (B) Quantification of mean CI-MPR intensity of control and patient’s fibroblasts was performed by comparing mean intensity from 30 cells per condition across 3 independent experiments (10 cells per condition per experiment). Mean intensity was averaged for each sample in each experiment and the ratio was statistically compared to a value of ‘1’ per experiment for a one-tailed paired t-test, p-value = 0.008. Notice that patient’s cells show significantly weaker staining with a more dispersed localization. (C) Patient’s and control fibroblasts were lysed, normalized by the Bradford protein assay and analyzed by SDS-PAGE and immunoblotting for endogenous CI-MPR. (D) Control and patient’s fibroblasts were immunostained with antibody to LAMP1 and co-stained with the actin marker phalloidin-555. We observed that ∼20% of cells exhibited abnormally large lysosomes, as shown in the figure. (E) Control and patient’s fibroblasts were lysed, normalized by Bradford protein assay and analyzed by SDS-PAGE and immunoblotting for LAMP1. (F) Quantification of fold differences in CI-MPR and LAMP1 protein levels in patient’s vs. normal fibroblasts from a minimum of three independent immunoblot assays per condition. Actin loading control was used to normalize the expression. Samples were compared with a one-tailed paired t-test, p-values: LAMP1 = 0.0163, CI-MPR = 0.0061. Notice that CI-MPR was less abundant and LAMP1 more abundant in the patient’s cells. In panels C and E, positions of molecular mass markers are indicated to the right. In A and D, nuclear staining by DAPI is shown in blue. CI-MPR and LAMP1 staining is shown in greyscale. In panel D, actin stained with phalloidin-555 is shown in red. Scale bars: 10 *μ*m. Error bars = standard deviation.

VPS51-KD and VPS52-KD HeLa cells^6,12^, as well as fibroblasts from VPS53-mutant PCCA2/PCH2E patients^23^, were previously found to have more numerous and larger lysosomes, as visualized by immunostaining with antibodies to the late-endosomal/lysosomal protein CD63. Immunostaining of the VPS51 mutant patient fibroblasts with an antibody to another lysosomal membrane protein, LAMP1, showed enlarged lysosomes in ∼20% of the cells (Fig. 6D). In addition, immunoblot analysis showed an increase in the levels of LAMP1 in patient vs. control fibroblasts (Fig. 6E,F).

## 3 Discussion

This study is the first to demonstrate the involvement of *VPS51* in a human disorder and provides further insight into the phenotypic effects resulting from disturbances in the GARP and EARP complexes.

### 3.1 Summary of the patient’s phenotype and comparison to pontocerebellar hypoplasia, PCCA2 and PEHO-like syndrome

The clinical phenotype observed in our patient is primarily characterized by neurodevelopmental manifestations, including severe global developmental delay, hypotonia, microcephaly, a seizure disorder, cortical vision impairment, and abnormal brain imaging with reduced size of the cerebellum and brainstem, hypoplasia of the corpus callosum, and cerebral white matter abnormalities. The generally static nature of the patient’s condition suggests that these findings may reflect primary developmental abnormalities of the brain. On the other hand, it is possible that her findings could be due to a progressive process, as described in patients with *VPS53* mutations^23,25^. Our patient’s normal early brain MRI scan followed by the discovery of multiple abnormalities on subsequent imaging would support the latter possibility. In either case, the similarities between our patient’s neurodevelopmental phenotype and that described in patients with *VPS53* mutations suggest that, in general, GARP/EARP dysfunction results in a severe neurodevelopmental disorder. Given that mutations in *VPS53* are classified as causing a specific form of pontocerebellar hypoplasia (PCH2E/PCCA2) and have also been associated with a PEHO-like syndrome, we propose that *VPS51* mutations may be responsible for a previously unclassified form of these disorders and that perturbation of GARP/EARP function has particularly severe effects on pontocerebellar development.

Our patient’s history of poor somatic growth, liver dysfunction, gastric volvulus, other health issues, and dysmorphic features is also notable and suggests that GARP/EARP dysfunction has systemic effects beyond the central nervous system. Interestingly, our patient had abnormal glycosylation testing without evidence of a specific congenital disorder of glycosylation (CDG), and her phenotype has significant overlap with that seen in CDGs. This suggests that GARP/EARP dysfunction may result in a complex phenotype that resembles CDGs both clinically and biochemically. The fact that she developed lower extremity edema at 6 years of age is also noteworthy in light of the previous description of limb edema in the two siblings with a PEHO-like syndrome caused by mutations in *VPS53* ^25^.

### 3.2 Neurological defects in animal models with mutations in GARP/EARP subunits

To date there have been no studies on the loss of VPS51 in a mammalian model. However, homozygous null mutations of the shared GARP/EARP subunits VPS52 or VPS53 in mouse cause embryonic lethality at E6.5 or E10.5-E11.5, respectively^16,18,19^. Likewise, homozygous null mutations of the GARP-specific VPS54 subunit in mouse are embryonic lethal at day E12.5^17^, with membrane blebbing into the lumen of the neural tube observed at days E10.5 and E11.5^19^. The earlier lethality resulting from *VPS52* and *VPS53* mutations relative to *VPS54* mutations suggests that combined loss of GARP and EARP has a more deleterious effect than that of GARP alone. Alternatively, EARP may be more important for viability. Distinguishing between these two possibilities will require the study of mice deficient in the EARP-specific VPS50 subunit.

The fact that the VPS51-deficient patient described here survived through embryonic development and infancy is consistent with the presence of residual levels of fully-assembled, functional GARP and EARP. This situation is analogous to that of *VPS54* mutations in the mouse, for which null mutations are embryonic lethal but the hypomorphic missense mutation in the *wr* mouse allows viability albeit with neurological defects^17^. The L967Q substitution in this mouse results in a protein that is conformationally unstable but partly incorporated into GARP^20^. The *wr* mouse was originally characterized phenotypically by an unsteady gait and head tremor, followed by progressive muscle weakness^28,29^. These defects were attributed to motor neuron degeneration similar to that in ALS. As in *wr*, the motor deficits in the patient described here may derive from loss of GARP function. Other findings in the patient could be due to partial loss of EARP function. In this regard, studies in *C. elegans* have implicated the EARP-specific VPS50 subunit in the formation of dense core vesicles, with downstream effects on locomotion^14,30^. Additionally, a mutation in TSSC1/EIPR-1, a protein associated with GARP/EARP^14,31^, has been shown to cause a similar phenotype in *C. elegans*^14^. These animal studies thus support the conclusion that the neurological deficits in the patient are due to partial loss of GARP and EARP.

### 3.3 Cellular effects of *VPS51* mutations

Characterization of skin fibroblasts from the patient revealed several abnormal phenotypes, including reduced GARP/EARP levels, increased levels of LAMP1, swelling of lysosomes and mislocalization of the CI-MPR. These phenotypes are consistent with the reduction in GARP/EARP levels causing lysosomal defects. The CI-MPR and related cation-dependent mannose 6-phosphate receptor (CD-MPR) mediate sorting of mannose 6-phosphate (M6P)-modified acid hydrolases from the TGN to endosomes^32^. The acidic pH of endosomes causes dissociation of the hydrolase-MPR complexes, after which the hydrolases traffic to lysosomes whereas the MPRs return to the TGN for further rounds of sorting. GARP promotes the tethering and fusion of the MPR carriers with the TGN^11^. Loss of GARP interrupts this cycle by preventing the retrieval of MPRs to the TGN, with consequent missorting of acid hydrolases towards the extracellular space^6,12^. The reduced levels of GARP thus likely explain the mislocalization of the CI-MPR in the patient cells. In this context, the swelling of lysosomes most probably results from accumulation of undegraded materials due to reduction of acid hydrolase levels, as previously observed upon knockdown of GARP subunits^6,12^. The increased levels of LAMP1 could result from decreased turnover of this protein in lysosomes or, alternatively, from increased synthesis due to activation of lysosome biogenesis pathways^33^.

The defect in MPR sorting in the patient’s cells is also likely to impact the Niemann-Pick C2 protein (NPC2), which, like the lysosomal acid hydrolases, is modified with M6P and sorted by the MPRs^27,34^. NPC2, together with NPC1, mediates export of cholesterol from the lumen of late endosomes and lysosomes to the cytosol or other organelles^35^. Consistent with this mechanism, cell lines with defects in GARP subunits, as well as the *wr* mouse, exhibit increased accumulation of cholesterol in lysosomes^20^. In addition, GARP mutations in yeast, including one analogous to that in human PCCA2, were shown to alter sphingolipid homeostasis^13^. It is thus possible that the patient could also have dysregulation of lipid metabolism. Little is known about other cargos that depend on GARP or EARP for recycling from endosomes. Further studies to identify such cargos will likely shed greater light into the pathogenesis of GARP/EARP deficiency.

### 3.4 Classification of *VPS51*-related disorders

PCCA2 caused by mutations in *VPS53* ^23^ has been proposed to be part of a more general disease classification with shared pathophysiology, termed the ‘Golgipathies’^36^. Golgipathies are inherited diseases that have a shared set of characteristics including microcephaly, white matter defects and intellectual disability. The causative proteins in this disease class are all functionally associated with the Golgi apparatus. Accordingly, the VPS51 mutations characterized here could be considered in the same class of diseases, occupying a subclass of “GARP/EARP deficiency disorders”. Further understanding of the phenotypic effects of mutations in VPS51 and other genes encoding components of the GARP/EARP complexes will require the identification and characterization of additional cases.

## 4 Materials and methods

### 4.1 Human subjects

The patient was seen for clinical genetic evaluations through the Greenwood Genetic Center. Parental informed consent for human subjects research was obtained through a protocol approved by the Institutional Review Board of Self Regional Healthcare (Greenwood, SC). De-identified skin fibroblasts from the patient and an unrelated control individual were shared with the National Institutes of Health laboratory for research purposes. For additional control fibroblasts (Fig. S2), unrelated anonymized fibroblasts from an unrelated project were used, as used previously^37^.

### 4.2 Whole-exome sequencing

DNA libraries were prepared from 3 *μ*g of genomic DNA isolated from the proband and parental peripheral blood samples, using the Agilent SureSelectXT Human All Exon v5 capture kit (Agilent Technologies, Santa Clara, CA). Briefly, DNA was fragmented using the Covaris S220 system (Covaris, Woburn, MA), and fragments of 150-200 bp were selected using AMPure XP beads (Beckman Coulter, Brea, CA). Fragments were subsequently end-repaired, adenylated at the 3’ end, ligated to sequencing adaptors, and then PCR-amplified using the SureSelectXT Library Preparation kit (Agilent Technologies). DNA was next purified using AMPure XP beads. The concentration of the enriched libraries was determined using a DNA-1000 chip on the 2100 Bioanalyzer (Agilent Technologies). 750 ng of each DNA library was used for hybridization and capture with the SureSelectXT Human All Exon v5 probes (Agilent Technologies). After hybridization, the RNA-bound DNA was retained, and unhybridized material was washed away. Captured fragments were amplified by PCR and purified. The quality of the enriched libraries was evaluated using a High-Sensitivity DNA chip on the 2100 Bioanalyzer. Libraries were quantified on a Qubit^XT^ 2.0 Fluorometer using a Qubit High Sensitivity kit (Life Technologies, Waltham, MA), and separate libraries were pooled and sequenced using an Illumina NextSeq 500 ^®^ Sequencing System (Illumina Inc., San Diego, CA) per the manufacturer’s protocol.

### 4.3 Bioinformatics

Exome sequencing generated a minimum of 97 million reads per patient sample with an average read length of 75 bp. Sequences were processed using NextGENe software (SoftGenetics, LLC, State College, PA) and trimmed for low-quality and duplicate reads. Reads were trimmed or rejected when the median Phred quality score was below 20 and/or had more than three bases with a Phred score *≤* 16. The sequences were then mapped to the February 2009 human reference assembly (GRCh37/hg19). Positions in the coding regions of the reference (with 25 bp flanking) were classified as variants if they had a read depth of >3 and a variant allele frequency of *≥* 12%. The variant reports were then converted into vcf files which were uploaded to Cartagenia Bench Lab NGS (Agilent Technologies, Santa Clara, CA) for filtering and subsequent analysis of variants of interest.

### 4.4 Exome data analysis

Variants were first matched to the Human Genome Mutation Database (HGMD); however, analysis of these variants did not yield promising gene candidates. Therefore, the unmatched variants were further analyzed. Benign variants were separated using the dbSNP, 1000 Genomes, ExAC and ESP6500 databases with an allele frequency cut off of 0.033. Next, quality and allele frequency filters were used to isolate variants with at least 20X read depth and 22% minor allele frequency. The resulting variants were then further analyzed using additional filters including those for splice site, functional effect prediction, and HPO phenotype. Reanalysis of the data was performed approximately 18 months after the initial analysis using the most current databases and literature available (ExAC, gnomAD, HGMD, PubMed).

### 4.5 Antibodies

The following antibodies were used for immunoblotting (IB) and/or immunofluorescence (IF): mouse anti-LAMP1 (H4A3; Developmental Studies Hybridoma Bank, Iowa City, IA), used at 1:1000 for IB and 1:500 for IF; mouse anti-CI-MPR (2G11; Abcam, Cambridge, United Kingdom), used at 1:500 for IF; homemade rabbit anti-CI-MPR, used at 1:1000 for IB; rabbit anti-VPS51 (HPA039650; Atlas Antibodies, Bromma, Sweden), used at 1:1000 for IB; homemade rabbit anti-VPS52^12^, used at 1:2000 for IB; rabbit anti-VPS53 (HPA024446; Atlas Antibodies), used at 1:1000 for IB; mouse anti-VPS50 (FLJ20097 monoclonal antibody M01, 2D11; Abnova, Taipei City, Taiwan), used at 1:1000 for IB; mouse anti-myc epitope (9E10; Santa Cruz Biotechnology, Santa Cruz, CA), used at 1:500 for IF; rat anti-HA epitope (3F10; Roche, Basel, Switzerland), used at 1:500 for IF; rabbit anti-GFP (A-11122; Thermo Fisher Scientific, Waltham, MA), used at 1:2000 for IB. HRP-conjugated secondary antibodies (Perkin Elmer, Waltham, MA) were used at 1:5000. Alexa Fluor-conjugated secondary antibodies for immunostaining (Thermo Fisher Scientific) were used at 1:1000. Alexa Fluor phalloidin-555 (Thermo Fisher Scientific) was used according to the manufacturer’s instructions.

### 4.6 DNA recombinant constructs

Plasmids encoding GFP fused to WT and mutant human VPS51 were constructed by Gibson assembly (New England Biolabs, Ipswich, MA) from two artificially synthesized dsDNA ‘gBlocks’ (Integrated DNA Technologies, Inc., Coralville, IA). The first gBlock (comprising nucleotides 1-1493) was codon-optimized for human codon usage to decrease DNA secondary structures. Synthetic gBlocks were assembled into pEGFP-C1 (Clontech, Takara Bio Inc., Kyoto, Japan) by Gibson assembly using a standard procedure (50°C incubation, 1 h). C-terminal 13myc epitope fusions were assembled into a modified pCI-neo (Promega, Madison, WI) by PCR amplification using VPS51-GFP-fusion templates and restriction digestion with EcoRI-SalI sites included in the primers (shared forward primer: 5’ CCTCGAGAATTCGCCACCATGGCCGCAGCCGCAGCGGCAGGG, WT and allele 2 reverse primer: 5’ CCCGGGGTCGACGCCGCGCTCGCAGATGACCTCAAC, allele 1 reverse primer: 5’ CCCGGGGTCGACAAACCCCGCCCCCGGAAAGGCCCG) and ligation into EcoRI-SalI digested vector. A cDNA encoding *Canis familiaris* Rab4B amplified from MDCK cells and cloned into pEGFP-C1 (Clontech) was a gift of Robert Lodge (NICHD, NIH). Human VPS54 cDNA was purchased (Origene, Rockville, MD) and subcloned into pCI-neo 13myc vector by Gibson assembly (fragment forward primer: 5’ TAGGCTAGCCCGAGAATTCGCCACCATGGC-TTCAAGCCACAGTTC; fragment reverse primer: 5’ TTAATTATCCCGGGGTCGACCCTCT-TCTGCTCCCAAATTTC; vector forward primer: 5’ GTCGACCCCGGGATAATTAAC; vector reverse primer: 5’ GAATTCTCGAGGCTAGCCTATAGTGAG). A plasmid encoding VPS50-13myc was previously described^8^. p3HA-N1 vector was generated by replacing the EGFP sequence of pEGFP-N1 with a 3HA sequence, followed by sub-cloning of codon-optimized human VPS51 into the HA and EGFP vector (for C-terminally tagged VPS51) using a PCR amplicon (forward primer: 5’ CCTCGAGAATTCGCCACCATGGCCGCAGCCGCAGCGGCAGGG; reverse primer: 5’ CGCGGTACCGTGCCGCGCTCGCAGATGACCT) digested with EcoRI-KpnI and ligated into an EcoRI-KpnI-digested vector backbone.

### 4.7 Cell culture and transfection

HeLa cells (ATCC, Manassas, VA) were cultured in ‘complete DMEM’ [DMEM supplemented with 10% fetal bovine serum (FBS), L-glutamine, penicillin-streptomycin (all from Corning, Corning, NY)], at 37°C, 5% CO_2_ and 95% relative humidity. HeLa cells were transfected with either Lipofectamine 2000 (Thermo Fisher Scientific) or Fugene 6 (Promega) according to the manufacturer’s instructions.

Skin biopsies obtained at the Greenwood Genetic Center were placed in sterile Petri dishes and minced using scalpels. Minced tissue was incubated for approximately 1 hour in collagenase solution (50 mg collagenase in 50 ml Minimum Essential Medium [MEM]) to dissociate cells from tissue fragments. The collagenase solution was removed and the remaining enzyme was de-activated using Fetal Bovine Serum (FBS). The cells were plated in ‘complete Chang Medium D’ (Chang D medium with 1% antibiotic-antimycotic solution) and allowed to grow in an incubator at 37°C, 5% CO_2_. After shipment to the NIH laboratory, human fibroblasts were cultured in Chang Medium D (Irvine Scientific, Santa Ana, CA) supplemented with MycoZap Plus-CL (Lonza, Walkersville, MD), at 37°C, 5% CO_2_ and 95% relative humidity. Human fibroblasts were transfected with Fugene HD (Promega) according to the manufacturer’s instructions.

### 4.8 Immunoprecipitation and biochemical analysis

Immunoprecipitation of GFP fusions was performed using magnetic GFP-trap beads (ChromoTek, Planegg-Martinsried, Germany) according to the manufacturer’s instructions. Briefly, cells transiently transfected on 100 mm cell culture plates were lysed for 30 min on ice in a buffer containing 10 mM Tris-HCl pH 7.4, 150 mM NaCl, 0.5 mM EDTA, 0.1% Nonidet P40 (NP40) and protease inhibitors (Roche, Basel, Switzerland). Lysates were centrifuged at 17,000 x g for 15 min at 4°C to remove cell debris. Cell extracts were combined with 25 *μ*l equilibrated GFP-Trap magnetic beads; volume was made up to 1 ml with lysis buffer without NP40. Incubation was for 2 h at 4°C under constant rotation. After magnetic separation to remove unbound material, beads were washed 3 times rotating at room temperature for 5 min with 1 ml of lysis buffer lacking NP40. Protein was eluted from beads by incubation in reducing Laemmli SDS-PAGE sample buffer (Bio-Rad) for 5 min 98°C. Samples were subsequently analyzed by SDS-PAGE and immunoblotting as described below.

For MG132 incubations, 800,000 HeLa cells were plated onto a 6-well dish and transfected with plasmids encoding VPS51-13myc or VPS51(Al1)-13myc. The following day, cells were split equally onto 4 wells each of a 12-well plate. The following day, a time-course (0, 2, 4 and 6 h) with 40 *μ*M MG132 (700 *μ*l per well) was performed, starting with a medium change at the 6 h time point to allow a simultaneous cell lysis across all samples. At the end of the time course, the cells were transferred to ice, washed in ice-cold PBS and lysed in 250 *μ*l lysis buffer [50 mM Tris pH 7.4, 150 mM NaCl, 1% Triton X-100 with protease inhibitors (Roche)]. Cells were incubated for 30 min on ice with pipetting every 10 min. After lysis, cells were transferred to an ice-cold 1.5 ml microcentrifuge tube, centrifuged at 17,000 x g for 15 min at 4°C to pellet cell debris, and the cell lysate was transferred to a new 1.5 ml microcentrifuge tube. Protein concentration was normalized across series by Bradford assay before IB probing.

Immunoblotting was performed by standard protocols using SDS-PAGE separation and subsequent transfer to PVDF membranes. Membranes were blocked for 1-2 h with 5% nonfat milk (Bio-Rad) in PBS-T [PBS supplemented with 0.05% Tween 20 (Sigma-Aldrich, St. Louis, MO)], and incubated overnight in primary antibody diluted in PBS-T with 3% BSA (Sigma-Aldrich) or PBS-T with 5% nonfat milk. Samples were washed 3 times for 5 min in PBS-T before being incubated for 1-2 h in HRP-conjugated secondary antibody (1:5000), diluted in PBS-T supplemented with 5% nonfat milk. Membranes were washed three times in PBS-T and once in PBS before being visualized using either Clarity western ECL blotting substrate (Bio-Rad) or SuperSignal West Femto maximum sensitivity substrate (Thermo Fisher Scientific).

### 4.9 Immunofluorescent staining

Cells for immunofluorescence microscopy (IF) were plated onto 12 mm cover glasses (EF15973A; Daigger, Vernon Hills, IL), transfected as described above, and fixed using 4% paraformaldehyde (Electron Microscopy Sciences, Hatfield, PA) in phosphate-buffered saline (PBS; KD Medical Columbia, MD). Cells were permeabilized at room temperature for 15 min in IF buffer [0.05% saponin (Sigma-Aldrich), 5% bovine serum albumin (BSA) (Sigma-Aldrich) in PBS]. Primary antibodies were diluted in IF buffer and incubated with cells for 1 h at room temperature. Cells were washed three times in IF buffer. Alexa Fluor secondary antibodies (Thermo Fisher Scientific) were diluted to 1:1000 in IF buffer and cells were incubated for 1 h at room temperature. Cells were washed 3 further times in IF buffer and once in water to remove PBS from the coverslip surface, and mounted to a glass cover slip in Immuno Mount [with or without 4,6-diamidino-2-phenylindole (DAPI); Electron Microscopy Sciences].

### 4.10 Microscopy

Imaging was performed on a Zeiss 710 or 880 microscope (Carl Zeiss, Oberkochen, Germany) with an oil-immersion 63X/1.40 NA Plan-Apochromat Oil DIC M27 objective lens (Carl Zeiss). Image settings (*i.e.*, gain, laser power, pinhole) were kept constant for images presented for comparison. ImageJ (http://imagej.nih.gov/ij/) was used for image processing, including cropping, average/maximum intensity projections and contrast and brightness adjustments, with any changes being kept consistent for comparable images. Images were assembled in Adobe Illustrator (Adobe Systems, San Jose, CA).

### 4.11 Quantification

Quantification of percentage of cells with VPS51-13myc at the TGN (Fig. 4C) was performed by counting a total of over 300 cells across 3 different independent experiments (over 100 cells per experiment per condition). Cells in which no TGN localization could be observed were classified in one category, and cells with any residual TGN localization were classified as having ‘VPS51-13myc at the TGN’. Percentages were calculated and compared with a one-tailed paired t-test.

Quantification of VPS51-positive Rab4B endosomes per cell (Fig. 4E) was calculated by manually counting VPS51-positive Rab4B endosomes in 62 cells (wt VPS51) and 64 cells (allele 2) from images taken across 3 independent experiments.

Quantification of fold differences in protein levels (Fig. 5 and 6) was done in ImageJ. Three independent immunoblot assays (i.e., different cell lysates) were performed per condition. Band intensity was calculated as mean intensity in ImageJ, with blot background removed by measuring mean intensity from an area without signal. For each sample, the corresponding actin loading control was used to normalize the expression. Samples were compared with a one-tailed paired t-test. Quantification of CI-MPR intensity of control and patient’s fibroblasts (Fig. 6) was analyzed in ImageJ by comparing mean intensity of 30 cells per condition across 3 independent experiments (10 cells per condition per experiment). Mean intensity was averaged for each sample in each experiment and the ratio was statistically compared to a value of ‘1’ per experiment for a one-tailed paired t-test.

Asterisks are indications of calculated statistical significance. ns: P > 0.05, * P *≤* 0.05, ** P *≤* 0.01, *** P *≤* 0.001, **** P *≤* 0.0001.

## 5 Declaration of Interests

The Greenwood Genetic Center receives revenue from diagnostic testing performed in the GGC Molecular Diagnostic Laboratory.

## 6 Acknowledgements

We thank the patient and her parents for their participation in this project and Robert Lodge for kind gifts or reagents. We also thank Melanie Jones, PhD for her initial analysis of the patient’s exome data, and Gail Stapleton, MS and Jessica Davis, MS for their assistance with the patient’s clinical evaluations. We would also like to thank Tal Keren-Kaplan, PhD and Amra Saríc, PhD for their critical reading of this manuscript. This work was funded by the Intramural Program of NICHD (project ZIA HD001607). Support was also provided to the Greenwood Genetic Center through the South Carolina Department of Disabilities and Special Needs.

## 8 Supplemental Data

### 8.1 Clinical Report

The patient is a 6-year-old Caucasian female who was first seen for a genetic evaluation at 55 days of age for failure to thrive and cholestatic hepatitis of unknown etiology. She was born to a healthy 30-year-old G2P1 mother via a cesarean delivery due to breech presentation following an uncomplicated pregnancy. She was discharged home with her mother and had a normal newborn screen but experienced early difficulties with weight gain and was hospitalized twice in the first 2 months of life for failure to thrive. Her family history was noncontributory, and her physical exam was primarily notable for weight, length, and head circumference all below the 3rd centile.

Her initial laboratory evaluation showed hyperbilirubinemia with a direct bilirubin of 2.7 mg/dL, elevated liver enzymes (GGT–8126 U/L, AST–8117 U/L, and ALT–8172 U/L), and normal alkaline phosphatase and creatine kinase levels. GALT enzyme analysis for galactosemia was normal and alpha-1 antitrypsin testing showed an MS protease inhibitor type. Urine organic acid analysis and plasma total and free carnitine levels, acylcarnitine profile, and amino acid analysis were non-diagnostic, with presence of N-acetyltyrosine but no other evidence of hepatorenal tyrosinemia. An eye exam showed no evidence of posterior embryotoxon to suggest Alagille syndrome, and sweat testing for cystic fibrosis was unsuccessful. A HIDA scan was normal and a liver biopsy showed mild giant cell changes, focal extra medullary hematopoiesis, reticuloendothelial and periportal iron, and electron microscopic findings of nonspecific neonatal giant cell hepatitis.

When reevaluated at almost 4 months of age, she was having ongoing difficulty with oral intake requiring tube feedings and concern for aspiration requiring thickened feedings. A neurological evaluation showed hypotonia. Her genetic test results included a negative 32 mutation screen for cystic fibrosis, a normal chromosome analysis at 650–850 band resolution, a genome wide microarray that showed a 139 kb deletion of chromosome 9p23 (later found to have been paternally inherited), and a normal a methylation analysis for Prader-Willi syndrome. Additional biochemical testing included a plasma 7-dehydrocholesterol level, plasma very long chain fatty acid analysis, and blood lactic and pyruvic acid levels, all of which were normal except for a mild elevation in lactic acid. Notably, however, serum transferrin isoelectric focusing showed a trace amount of hypoglycosylation manifested by a slightly low percentage of trisialo-transferrin and slightly increased percentages of asialo-and monosialo-transferrin. This testing was repeated and showed similar findings, with persistent mild increases in the latter two compounds, but a subsequent reanalysis was normal and a carbohydrate transferrin profile showed a normal mono-/di-oligosaccharide transferrin ratio and a normal a-oligo-/di-oligo-transferrin ratio.

Thereafter, she had ongoing feeding problems requiring placement of a gastrostomy tube and was also incidentally found to have an organo-axial gastric volvulus. Due to persistent liver dysfunction, a liver biopsy was repeated and showed mild portal chronic inflammation and focal bridging hepatic fibrosis. She also required ear tube placement and a hospitalization for RSV. An ophthalmological exam showed delayed visual maturation, and an EEG and brain MRI were normal. When reevaluated at almost 10 months of age, she had global developmental delay with an inability to sit independently and was receiving multiple therapies. Her weight and length were below the 3rd centile, and her head circumference was markedly below the 3rd centile. Her exam was notable for brachycephaly with a depression in the posterior skull, prominent epicanthal folds, an irregular right ear helix, an upturned nasal tip, an open mouth with a hypotonic facial appearance, a rounded face with full cheeks, an enlarged liver, and marked truncal hypotonia with a head lag. A skull X-ray was obtained and was normal.

At age 12 months, repeat plasma amino acid and urine organic acid analyses as well as POLG1 sequencing were normal. She was hospitalized at 15 months for a pleural effusion. She was subsequently evaluated at an outside institution and was found to have an abnormal pattern of N-and O-glycosylation, which was also seen on repeat testing. Because of concern for a congenital disorder of glycosylation (CDG) predisposing to coagulopathy, an antithrombin III level was checked and was slightly low (78%; normal anterior palate, full-120) while a PT, PTT, and INR were normal. At age 23 months, she had an ALT of 181, an AST of 318, and an alkaline phosphatase of 328, and a CBC showed a low white cell count of 4,500 with an absolute neutrophil count of 900 cells/mm^3^. Her eye exam was notable for cortical vision impairment with an estimated visual acuity of 20/400, intermittent esotropia, and normal optic nerves and retinas. An auditory brainstem response showed slight unilateral hearing loss. EEG monitoring showed an increased risk of seizures without overt seizures, and she was placed on levetiracetam shortly after turning 2 years of age.

When seen again at age 2 years 4 months, she was taking ursodiol and being treated for gastroesophageal reflux. She was making extremely slow developmental progress despite continued therapies, in that she could roll from side to side but not all the way over, was unable to sit independently, and could not say any words. All of her growth parameters remained below the 3rd centile with marked microcephaly, a narrow forehead, epicanthal folds, long eyelashes, intermittent exotropia and nystagmus, slightly overfolded ears, an upturned nasal tip, a thin upper lip, a high/narrow anterior palate, full/rounded cheeks, a low posterior hairline, mild liver enlargement, hypoplastic labia minora, mild stiffness in the knees, single flexion creases on the fifth fingers, mild clubbing of the thumbnails and index fingernails, increased hair on the upper back, and marked hypotonia with a continued head lag. Whole exome sequencing had previously been initiated but was not yet complete.

By age 2 years 11 months, she had required a brief ICU stay for RSV and had been diagnosed with sleep apnea. She was receiving all nutrition through her gastrostomy tube. A repeat EEG had shown stable abnormalities, and an eye exam showed persistent cortical vision impairment. Developmentally, she was still not using words or making significant progress with her motor skills despite ongoing therapies. Her physical exam was essentially unchanged. The whole exome sequencing had been completed by this time.

She was subsequently hospitalized for an episode of adenoviral pneumonia with low platelet and white cell counts, and her gastroenterologist had noted ongoing hepatitis. A routine EEG showed significant seizure activity despite absence of clinical seizures, prompting an overnight EEG that showed possible electrical status epilepticus of sleep (ESES).

When seen again at age 4 years 3 months, her medications included levetiracetam, ursodiol, ranitidine, and clonidine. She remained G-tube dependent and had severe constipation with an episode of fecal impaction prompting the use of polyethylene glycol. She had developed warts on her hand due to a suspected HPV infection, and her dentist was concerned about thin enamel on her posterior molars. She had ongoing mild sleep apnea and had been found to have iron deficiency prompting treatment with ferrous sulfate. She had intermittent esotropia with some improvement in her visual acuity (20/200). She was able to roll into a prone position but could not hold her head up and was not saying any words. Her growth parameters remained below the 3rd centile with marked microcephaly, and her exam findings were otherwise unchanged except for the presence of frequent body movements.

A subsequent repeat brain MRI at 4 years 6 months showed hypoplasia of the pons, cerebellar vermis, and corpus callosum, possible hypoplasia of the hippocampus, a suspected Dandy-Walker variant with the 4th ventricle opening into a prominent infra-vermian cistern, and symmetric signal abnormalities in the periventricular white matter. Per the neurologist’s reading, it was not clear whether the abnormalities of the pons and cerebellum represented hypoplasia or atrophy, given the lack of an interval MRI between this study and the one done much earlier in life.

By age 5 years 3 months, she was having persistent abnormal EEG activity due to ESES that required treatment with diazepam and valproic acid. She had again been hospitalized for RSV and on another occasion for the flu. She continued on ursodiol for her liver dysfunction, and an elevated copper level had been identified. Due to concern for aspiration of oral secretions, a salivary gland procedure was planned. She continued to have cortical vision impairment, hypotonia, sleep apnea, and constipation. An orthopedic evaluation had shown partial dislocation of one hip, and her dentist had noted oddly shaped molars. She was not making any major developmental gains despite ongoing therapies. Her exam findings were essentially unchanged, with a weight and height below the 3rd centile and a head circumference at the 50th centile for age 7 months. An exome reanalysis was requested at that time. Metabolomic testing was also done and showed a number of abnormal small metabolite levels in plasma but no evidence of a perturbation in a specific metabolic pathway.

When last seen at age 6 years 3 months, she had recently had another brain MRI that showed generalized brain atrophy with disproportionate cerebellar atrophy, near absence of the cerebellar vermis with an enlarged cisterna magna reflecting a possible Dandy-Walker variant, loss of brainstem volume, bilateral atrophy of the hippocampi, and loss of cerebral white matter volume with a markedly small corpus callosum and generalized hyperintense T2 signal. These were felt to reflect chronic abnormalities that were unchanged from the previous study. Her seizures were more difficult to control and she had not made significant developmental gains. She had developed chronic diarrhea and had required hospitalization for a UTI. She had also developed asymmetry of her lower extremities with associated pitting edema for which further imaging was planned.

### 8.2 Supplemental figures

**Figure S1:**
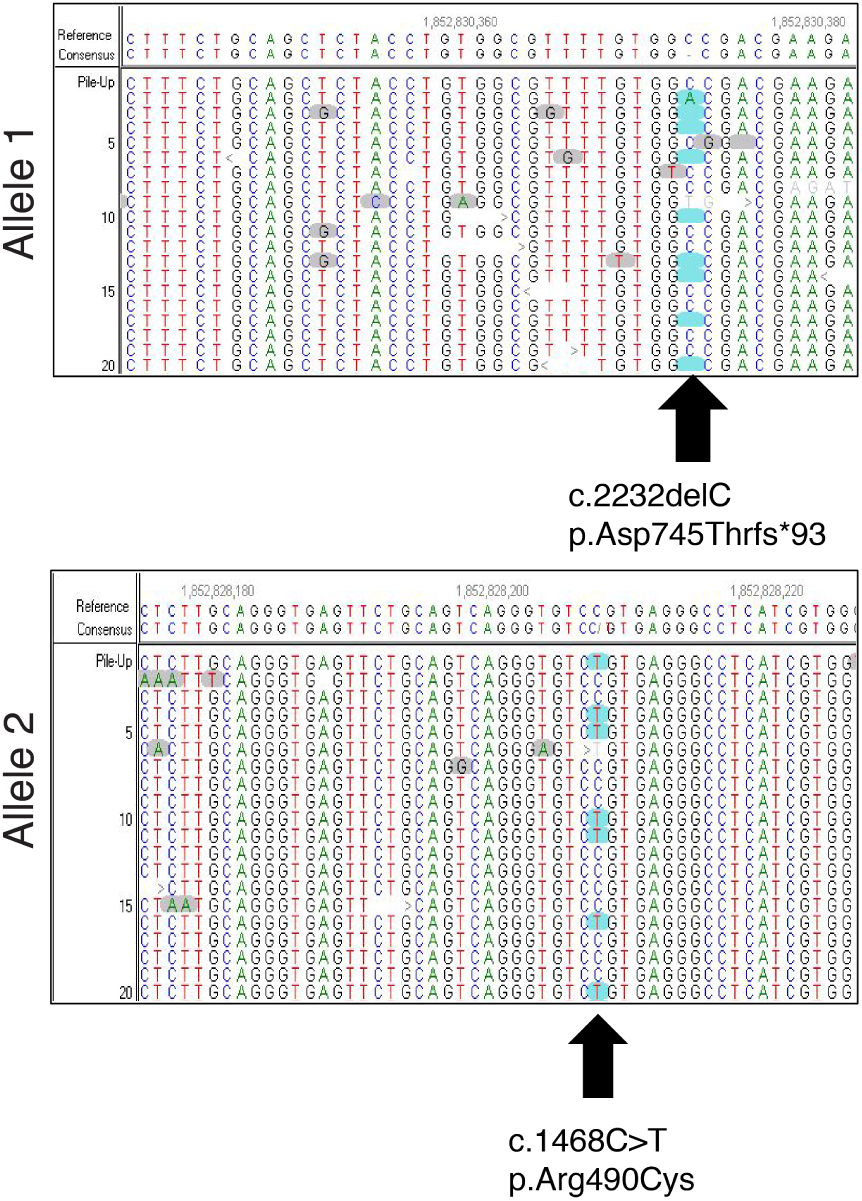
Identification of mutations in *VPS51* by whole-exome sequencing.

**Figure S2:**
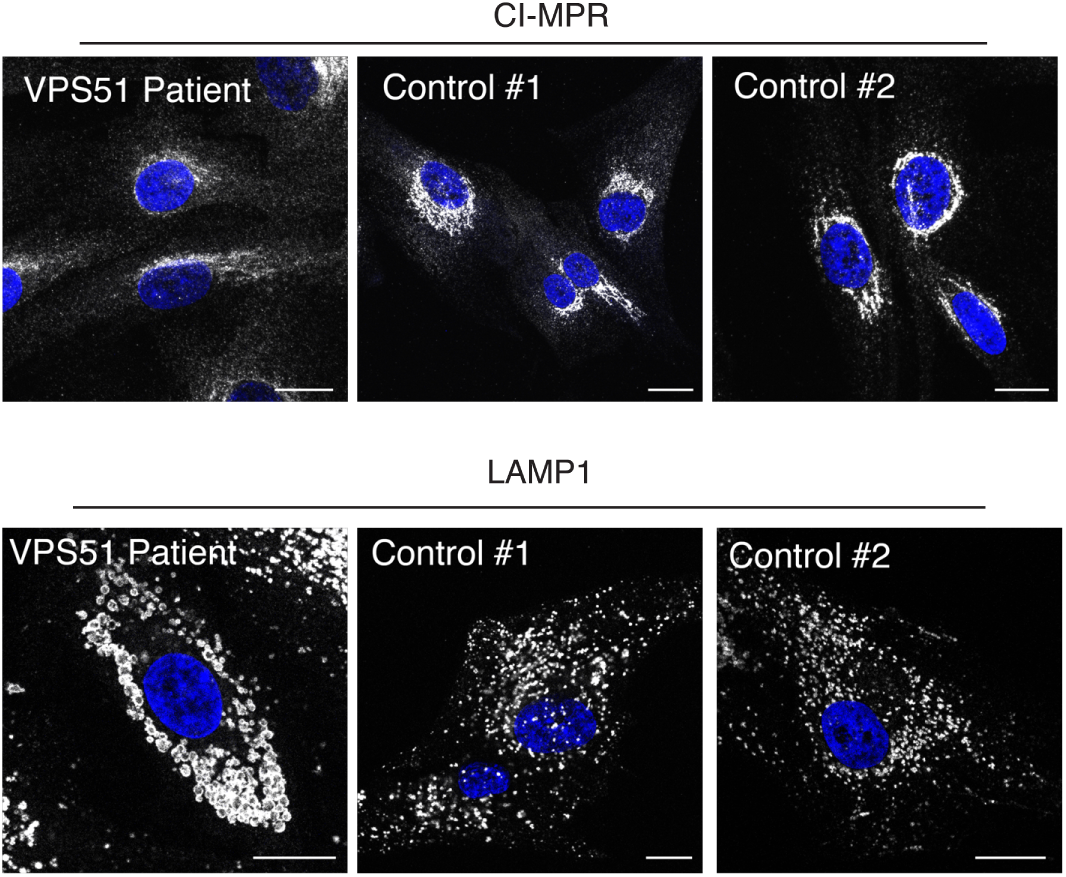
Comparison of CI-MPR and LAMP1 staining in patient’s fibroblasts and two control fibroblast lines. *VPS51* patient’s fibroblasts and fibroblasts from two independent human controls were immunostained with antibody to either CI-MPR or LAMP1. Left panel is patient cells, middle panel is control cells used elsewhere in manuscript, right panel is an additional control line from an independent project in the lab^37^. Nuclear staining by DAPI is shown in blue, single channels in greyscale. Scale bars: 10 *μ*m.

